# Enrichment in conservative amino acid changes among fixed and standing missense variations in slow evolving proteins

**DOI:** 10.1101/644666

**Authors:** Mingrui Wang, Dapeng Wang, Jun Yu, Shi Huang

## Abstract

Proteins were first used in the early 1960s to discover the molecular clock dating method and remain in common usage today in phylogenetic inferences based on neutral variations. To avoid substitution saturation, it is necessary to use slow evolving genes. However, it remains unclear whether fixed and standing missense changes in such genes may qualify as neutral. Here, based on the evolutionary rates as inferred from identity scores between orthologs in human and Macaca monkey, we found that the fraction of conservative amino acid mismatches between species was significantly higher in slow evolving proteins. We also examined the single nucleotide polymorphisms (SNPs) by using the 1000 genomes project data and found that missense SNPs in slow evolving proteins also had higher fraction of conservative changes, especially for common SNPs, consistent with more natural selection for SNPs, particularly rare ones, in fast evolving proteins. These results suggest that fixed and standing missense variations in slow evolving proteins are more likely to be neutral and hence better qualified for use in phylogenetic inferences.

## Introduction

Inferring phylogenetic relationships of species is an important and interesting area of evolutionary research. Before the molecular era, such relationships were built by mostly using comparative data from phenotypic features and fossils. Since the early 1960s, protein and later DNA sequence comparisons became increasingly dominant in building evolutionary trees (Zuckerkandl, Pauling 1962; Margoliash 1963; Doolittle, Blombaeck 1964; Fitch, Margoliash 1967). A seeming relationship between protein non-identity and time of separation had led to the molecular clock hypothesis, which assumes a constant and similar evolutionary rate among species (Zuckerkandl, Pauling 1962; Margoliash 1963; Kumar 2005). Thus, gene non-identity between species is thought to be largely a function of time. The molecular clock has in turn led Kimura to propose the neutral theory to explain nature: sequence differences between species are largely due to neutral changes (not adaptive evolution) (Kimura et al.1998). However, the molecular clock assumption may be unrealistic (Ayala 1999; Pulquerio, Nichols 2007). The neutral theory remains an incomplete explanatory theory as it predicts a constant substitution rate as measured in generations whereas the observed molecular clock is measured in years(Hu et al. 2013; Kern, Hahn 2018).

The initial justification for using proteins to build phylogenetic trees is the observation of a seemingly linear relationship between pairwise distances (or mismatches) and time of separation as determined by the fossil records (Zuckerkandl, Pauling 1962; Margoliash 1963; Doolittle, Blombaeck 1964; Fitch, Margoliash 1967), which has led to the distance matrix method (Fitch, Margoliash 1967). The neutral theory further provides an explanation for such a relationship that the substitution rate under selective neutrality is expected to be equal to the mutation rate (Kimura 1983). If however, mutations/substitutions are not neutral or under natural selection, the substitution rate would be affected by the population size and the selection coefficient, which are unlikely to be constant among the lineages. Thus, the distance matrix methods are sound provided that one uses neutral variants that accumulate to increase genetic distances in a nearly linear fashion common to the species concerned.

In phylogenetic inferences, one well known factor that may increase the uncertainty of the analysis is substitution saturation (Steel, Lockhart, Penny 1993; Philippe, Forterre 1999; Huang 2010; Brandley et al. 2011; Huang 2012; Soubrier et al. 2012). In saturation, the amount of substitutions or genetic distances would cease to be linearly related to time. Intuitively, the issue of saturation can be easily solved by using slow evolving genes as defined by high identity among species. However, such genes are well known to be under stronger purifying selection as defined by dN/dS ratio, which means that a new mutation has lower probability of being fixed (Cai, Petrov 2010). However, purifying selection as detected by the dN/dS method is largely concerned with non-observed mutations and says little about the fixed or observed variations. And phylogenetic inferences are all concerned with observed variants. Nonetheless, it remains to be determined whether fixed and standing missense substitutions in slow evolving genes may qualify as neutral.

We here found that the proportion of conservative mismatches between species was higher in the slowest evolving set of proteins than in fast evolving proteins. Using datasets from the 1000 genomes (1KG) project phase 3 dataset (Auton et al. 2015), we also found that missense single nucleotide polymorphisms (SNPs) from the slowest evolving set of proteins, especially those with high minor allele frequency (MAF), were enriched with conservative amino acid changes, consistent with such changes being under less natural selection. We suggest that the distance matrix methods, together with the new method here for selecting slow evolving sequences or neutral variants, may be best qualified for inferring realistic evolutionary trees.

## Methods

### Classification of proteins as slow and fast evolving

The identification of slow evolving proteins and their associated SNPs were as previously described (Yuan et al. 2017). Briefly, we collected the whole genome protein data of Homo sapiens (version 36.3) and Macaca mulatta (version 1) from the NCBI ftp site and then compared the human protein to the monkey protein using local BLASTP program at a cut-off of 1E-10. We only retained one human protein with multiple isoforms and chose the monkey protein with the most significant E-value as the orthologous counterpart of each human protein. The aligned proteins were ranked by percentage identities. Proteins that show the highest identity between human and monkey were included in the set of slow or the slowest evolving (including 423 genes 003E 304 amino acid in length with 100% identity and 178 genes > 1102 amino acid in length with 99% identity between monkey and human). The rest are all considered as fast evolving proteins. The cut off criterion was based on the empirical observation of low substitution saturation, and the finding that missense SNPs from the slow set of proteins produced genetic diversity patterns that were distinct from those from the fast set (Yuan et al. 2017).

### SNP selection

We downloaded the 1KG phase 3 data and assigned SNP categories using ANNOVAR (Auton et al. 2015). We then picked out the missense SNPs located in the slow evolving set of genes from the downloaded VCF files (Yuan et al. 2017). MAF was derived from AF (alternative allele frequency) values from the VCF files. Missense SNPs in fast evolving genes included all those from 1KG that are not from the slow evolving set.

### Scoring conservative amino acid replacements

For fixed substitutions as revealed by BLASTP, conservative changes were scored as those that are counted as “positive” mismatches in BLASTP alignment, which was largely based on having positive values in the BLOSUM62 matrix (Pearson 2013). The degree of physical/chemical change in an amino acid missense mutation was ranked by a scoring series, −3,-2, -1, 0, 1, 2, 3, in the BLOSUM62 matrix with more positive values representing more conservative changes. For missense SNPs, we assigned each amino acid mutation a score based on the BLOSUM62 matrix.

## Results

### Fixed amino acid mismatches and evolutionary rates of proteins

We determined the evolutionary rates of proteins in the human proteome by the percent mismatches between the human proteins and their orthologs in Macaca monkey. As described previously, the slow set contains the slowest evolving 423 genes (> 304 amino acid in length) with 100% identity and 178 genes (> 1102 amino acid in length) with 99% identity between monkey and human (Yuan et al. 2017). This group was identified based on the observation that missense SNPs located in these proteins produced distinct genetic diversity patterns from those by missense SNPs in faster evolving proteins or those by a randomly selected set consisting of mostly noncoding SNPs (Yuan et al. 2017). We then divided the proteins into several groups of different evolutionary rates and compared the proportion of conservative amino acid mismatches in each group. Conservative changes were scored as those that are counted as “positive” mismatches in BLASTP alignment (>0 based on scale found in the BLOSUM62 matrix) (Pearson 2013).

The mismatches between two species would have one of the two residues or alleles as ancestral in the case of slow evolving proteins yet to reach saturaton (no independent mutations occurring at the same site among species) and so a mismatch due conservative changes would involve a conservative mutation during evolution from the ancestor to extant species. But at saturation for fast evolving proteins where a site may encounter multiple mutations, while a drastic mismatch would necessarily involve a drastic mutation, it is possible for a conservative mismatch to result from at least two independent drastic mutations (If the common ancestor has Arg at some site, a drastic mutation event at this site occurring in each of the two species, Arg to Leu in one and Arg to Ile in the other, may lead to a conservative mismatch of Leu and Ile).

Thus, a conservative mismatch at saturation just means less physical/chemical differences between the two concerned species and says little about the actual mutation events. Lower fraction of conservative mismatches at saturation for fast evolving proteins would mean more physical/chemical differences between the two species, which may more easily translate into functional differences for natural selection to see and act upon.

To verify that the slowest evolving proteins with length >1102 amino acids included in the slow set is distinct from the fast set, we first compared proteins with length >1102 amino acids with no gaps in alignment (Table 1, Figure 1A) or with gaps (Table 1, Figure 1B). There was a general correlation between slower evolutionary rates and higher fraction of conservative changes, with a significant drop in the fraction of conservative changes between the slowest evolving, which was included in the slow set that has monkey-human identity > 99%, and the next set (Figure 1).

**Table 1.**
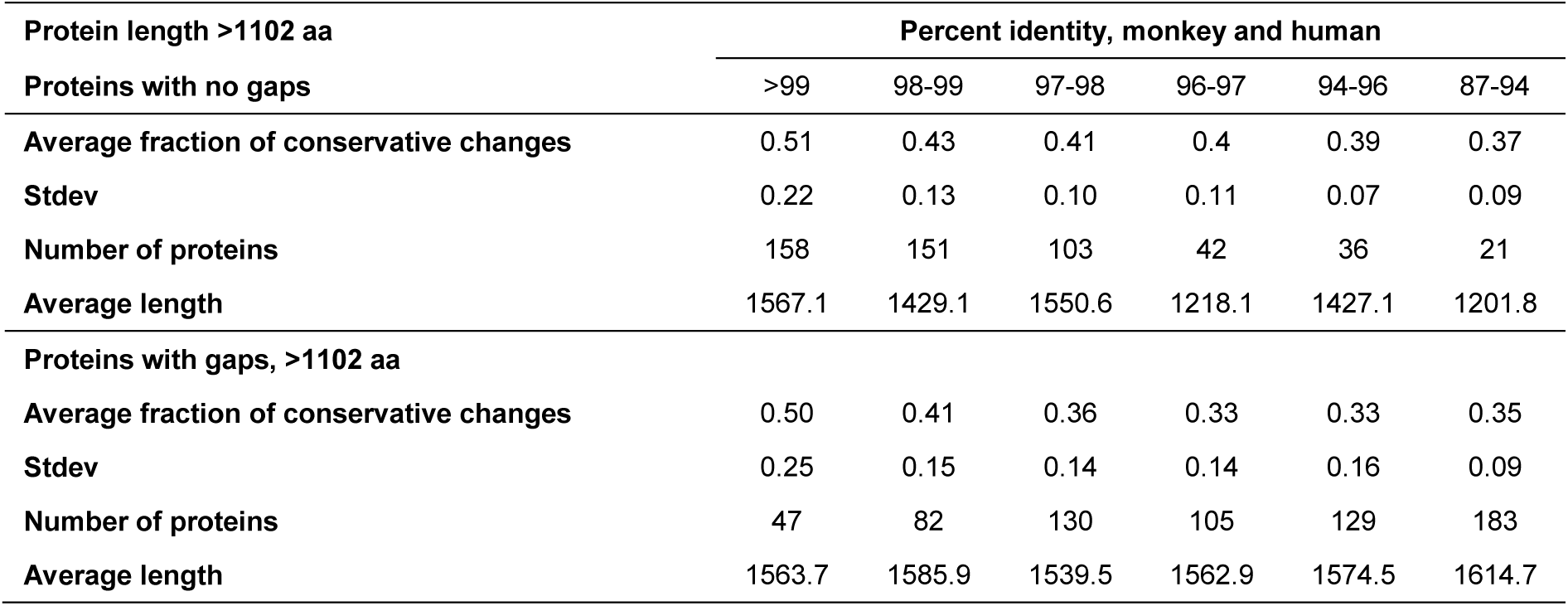
Conservative mismatches in proteins with length > 1102 amino acids.

| Protein length >1102 aa | Percent identity, monkey and human |  |  |  |  |  |
| --- | --- | --- | --- | --- | --- | --- |
|  | >99 | 98-99 | 97-98 | 96-97 | 94-96 | 87-94 |
| **Proteins with no gaps** |  |  |  |  |  |  |
| Average fraction of conservative changes | 0.51 | 0.43 | 0.41 | 0.4 | 0.39 | 0.37 |
| Stdev | 0.22 | 0.13 | 0.10 | 0.11 | 0.07 | 0.09 |
| Number of proteins | 158 | 151 | 103 | 42 | 36 | 21 |
| Average length | 1567.1 | 1429.1 | 1550.6 | 1218.1 | 1427.1 | 1201.8 |
| **Proteins with gaps, >1102 aa** |  |  |  |  |  |  |
| Average fraction of conservative changes | 0.50 | 0.41 | 0.36 | 0.33 | 0.33 | 0.35 |
| Stdev | 0.25 | 0.15 | 0.14 | 0.14 | 0.16 | 0.09 |
| Number of proteins | 47 | 82 | 130 | 105 | 129 | 183 |
| Average length | 1563.7 | 1585.9 | 1539.5 | 1562.9 | 1574.5 | 1614.7 |

**Figure 1.**
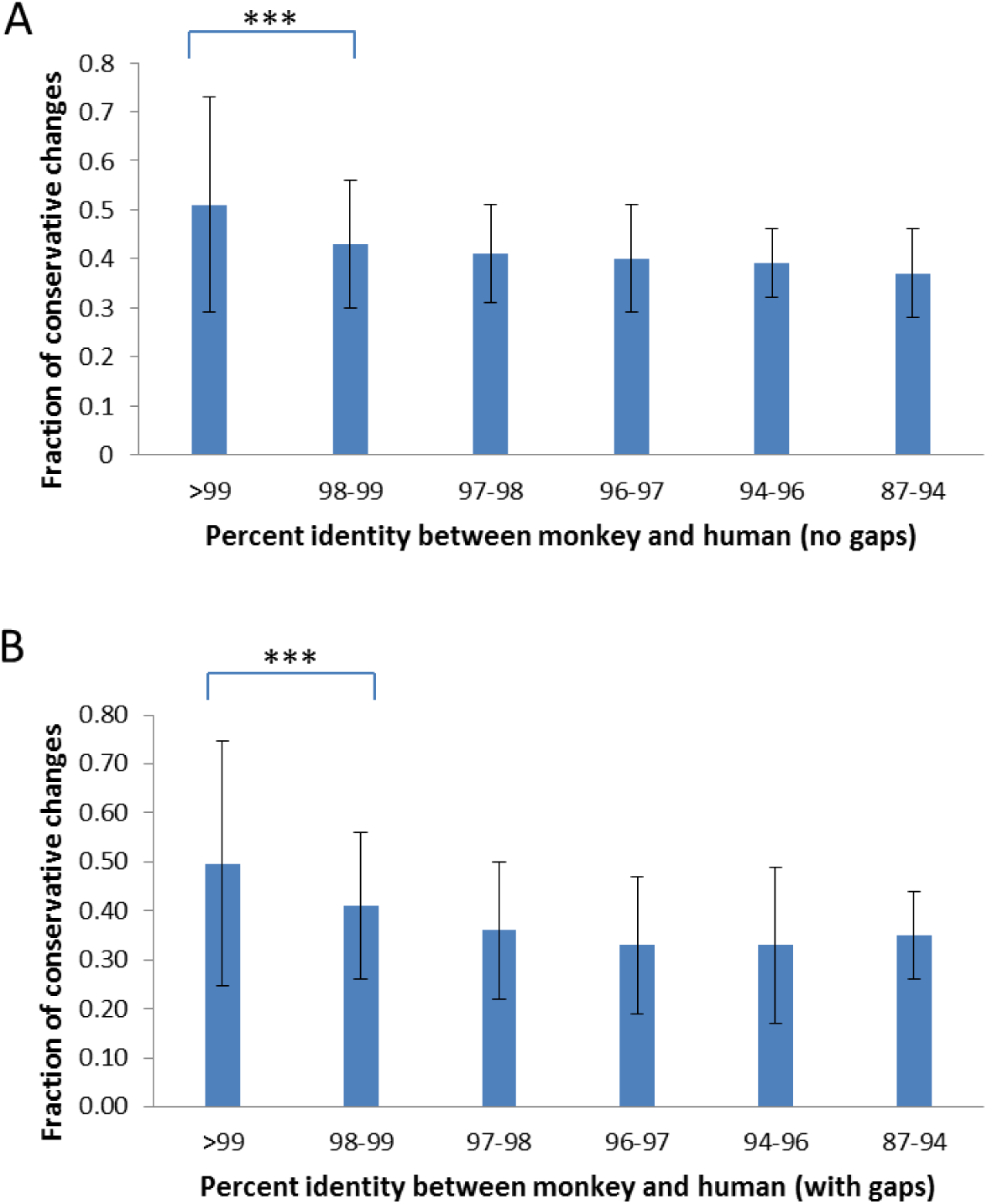
Fixed conservative amino acid mismatches in proteins of different evolutionary rates with length > 1102 amino acids. Shown are fractions of conservative changes in proteins with no gaps in alignment **(A)** and with gaps **(B)**. ***: P<0.001.

Proteins with alignment gaps showed slightly lower fraction of conservative changes than those without gaps, consistent with lower sequence conservation (hence faster evolving) in proteins with alignment gaps. We further studied the remaining proteins with shorter protein length (200-1102 amino acids). The set with no gaps showed proportions of conservative changes (0.44-0.38) similar to that of the fast sets of longer proteins (>1102 aa) with no gaps (0.43-0.37, Table 2 and Figure 2). The set with gaps showed similar results as that of the fast sets of longer proteins with gaps (0.41-0.33 vs 0.41-0.33). Thus, other than the slow set having a substantially larger fraction of conservative changes (0.51-0.50), all other proteins showed similarly low levels of conservative changes (0.44-0.33). The results confirmed the distinctive nature of the slow set of proteins as initially found by using missense SNPs in these proteins (Yuan et al. 2017). As fast evolving proteins are at saturation, low levels of conservative changes appear to be a feature of saturation.

**Table 2.** Conservative mismatches in proteins with length 200-1102 amino acids.

| Protein length 200-1102 aa | Percent identity, monkey and human |  |  |  |
| --- | --- | --- | --- | --- |
|  | 95-99 | 90-95 | 80-90 | 60-80 |
| **Proteins with no gaps** |  |  |  |  |
| Average fraction of conservative changes | 0.44 | 0.39 | 0.38 | 0.39 |
| Stdev | 0.26 | 0.12 | 0.11 | 0.08 |
| Number of proteins | 6728 | 1254 | 244 | 19 |
| Average length | 485.1 | 414.9 | 341.4 | 362.5 |
| **Proteins with gaps** |  |  |  |  |
| Average fraction of conservative changes | 0.37 | 0.34 | 0.33 | 0.30 |
| Stdev | 0.21 | 0.13 | 0.13 | 0.12 |
| Number of proteins | 1389 | 1008 | 647 | 200 |
| Average length | 593.7 | 554.1 | 464.3 | 451.9 |

**Figure 2.**
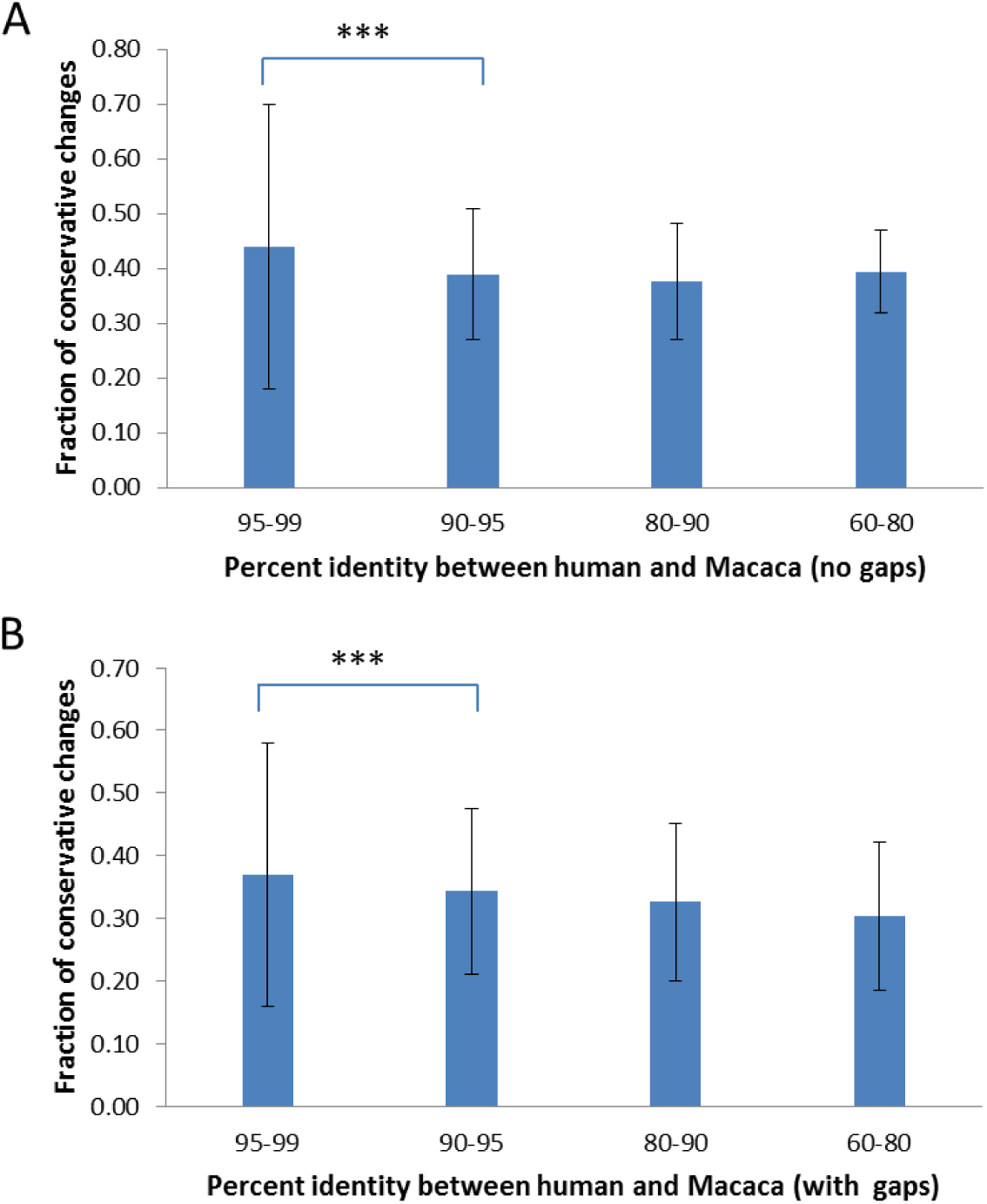
Fixed conservative amino acid mismatches in proteins of different evolutionary rates with length 200-1102 amino acids. Shown are fractions of conservative changes in proteins with no gaps in alignment **(A)** and with gaps **(B)**. ***: P003C0.001.

### Standing amino acid variants and evolutionary rates of proteins

We next studied missense SNPs found in proteins of different evolutionary rates by using 1KG dataset (Auton et al. 2015). There were 15271 missense SNPs from the slow evolving set of proteins (>1102aa with 99% identity and > 304 aa with 100% identity) and 546297 missense SNPs in fast set (all proteins that remain after excluding the slow set). We assigned each amino acid change found in a missense SNP a conservation score (−3, −2, −1, 0, 1, 2, or 3) as described in the BLOSUM62 matrix with higher values representing more conservative changes in the physical/chemical properties of the amino acids (Pearson 2013). The amount of SNPs in each score category was then enumerated. We found that missense SNPs in the slow evolving set of proteins had lower fraction of drastic mutations (with score −3, and-2) and higher fraction of conservative mutations (with scores 3, 2, and 1) relative to those in the fast evolving set of proteins (P<0.001 for all score categories except for −1 and 0, Figure 3). The fraction of conservative mutations (with scores >0) in the slow evolving set was significantly higher than that of fast set (P<0.001, Figure 3).

**Figure 3.**
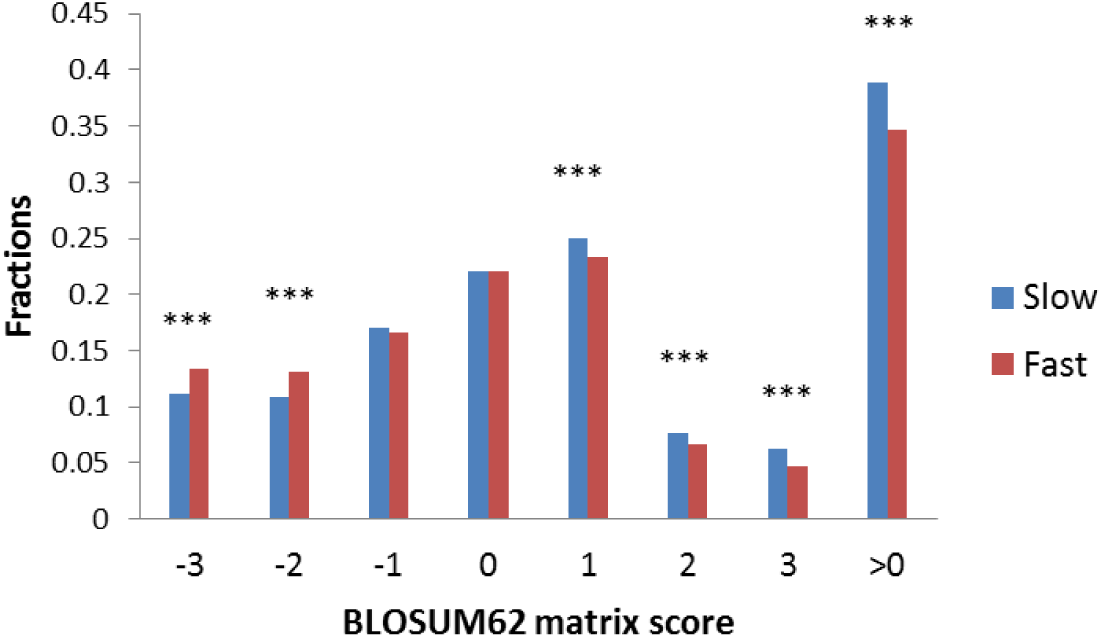
Conservative changes in standing missense substitutions in proteins of different evolutionary rates. Missense SNPs from either slow or fast group of proteins were classified based on the scores in the BLOSUM62 matrix. Shown are the fractions of each class. ***, P<0.001.

To test for neutrality or natural selection regarding conservative changes, we next divided the slow evolving set of missense SNPs into three groups of different minor allele frequency (MAF) as measured in Africans (similar results were found for other racial groups). For fast evolving proteins at saturation, low MAF values of a missense SNP would mean more purifying selection and so SNPs with low MAF are expected to have lower proportion of conservative amino acid changes as such changes may mean too little functional alteration to be under natural selection. The results showed that, for missense SNPs in the fast evolving set of proteins, the common SNPs with MAF 003E0.05 showed higher fraction of conservative changes than the rare SNPs with MAF<0.005 (P<0.001), indicating more natural selection for the rare SNPs in the fast set (Figure 4). While SNPs in the fast set showed similar fractions of conservative changes across groups, there was a trend of higher proportion of conservative changes as MAF values increase from >0.005 to >0.01 to >0.05 for SNPs in the slow set, consistent with less natural selection for common SNPs in the slow set (Figure 4). Each of the three groups based on MAF values in the fast set showed significantly lower fraction of conservative changes than the respective group in the slow set (P<0.001), indicating more natural selection for SNPs in the fast set (Figure 4). The results indicate that common SNPs in slow evolving proteins had more conservative changes that were under less natural selection.

**Figure 4.**
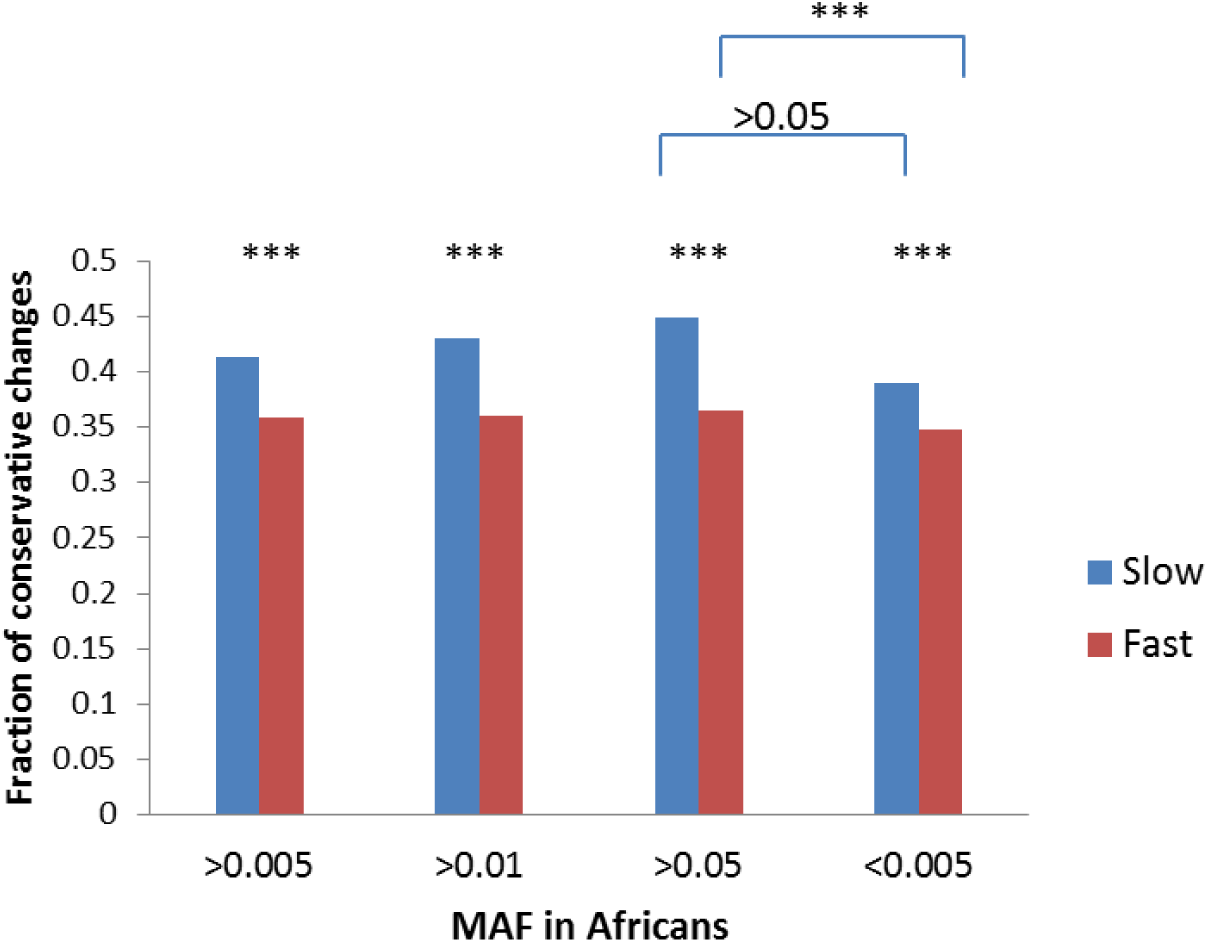
Conservative changes in missense SNPs with different MAF. SNPs from either fast or slow evolving proteins were classified based on MAF values and the fraction of conservative changes in each class is shown. ***, P<0.001.

## Discussion

Our results here showed that, contrary to naïve expectations, fixed and standing changes in slow evolving proteins were enriched with conservative amino acid mismatches. Furthermore, such conservative changes are also more neutral or under less natural selection.

Fixed and stranding conservative variants in slow evolving proteins may be under less natural selection for several reasons. First, conservative changes may not alter protein structure and function as dramatically as the drastic changes, which may make it harder for natural selection to see or act.

Second, as the lower fraction of conservative changes in SNPs in fast evolving proteins at saturation can only be explained by natural selection, the much higher fraction of such changes in high MAF SNPs in slow evolving proteins cannot be due to selection. For fast evolving proteins that have reached mutation saturation, a variant site within such proteins would have encountered multiple mutations. In contrast, for slow evolving proteins that are still at the linear phase of accumulating mutations, a mutant site would have chance for only one mutation to occur. An examination of the codon code reveals that, while conservative changes largely take just one nucleotide mutation, drastic changes often take 2 or 3 mutations at 2 or 3 sites. Thus, drastic changes is expected to be a function of mutation saturation, which is exactly what our results of lower fraction of fixed conservative changes in fast evolving proteins had shown.

Thirdly, as fixed variants cannot be fixed because of negative selection, they are either neutral or under positive selection. Indeed, fast evolving proteins are known to be under more positive selection (Cai, Petrov 2010; Yuan et al. 2017), which implies that fixed variants in slow evolving proteins can only be more neutral. Even if slightly deleterious mutations are fixed, it would not be because of selection but rather because of random drift. It makes sense for slow evolving proteins to be spared by positive selection because a mutation that takes a long time to arrive would be useless for quick adaptive needs.

Finally, SNPs in the slow set may be under purifying selection if they are drastic changes or no selection if they are conservative changes (assuming no positive selection as explained above). Thus, one would expect less conservative changes in the rare SNPs in the slow set, much more so than SNPs in the fast set under saturation. Our results are consistent with such expectations.

It is easy to tell optimum/maximum saturation genetic distances from linear distances as described previously (Huang 2010). Imagine a 100 amino acid protein with only 1 neutral site. In a multispecies alignment involving at least 3 species, if one finds only one species with a mutation at this neutral site while all other species have the same non mutated residue, there is no saturation. However, if one finds that nearly every species has a unique amino acid, one would conclude mutation saturation as there are multiple independent substitution events among different species at the same site and repeated mutations at the same site do not increase distance. We have termed those sites with repeated mutations “overlap” sites (Huang 2010). So, a diagnostic criterion for saturated maximum distance is the proportion of overlap sites among mismatched sites between two species. Saturation would typically have 50-60% overlapped sites that are 2-3 fold higher than that expected under no-saturation (Huang 2010; Luo, Huang 2016). It is not expected to have near 100% overlapped sites because certain sites may only accommodate 2 or very few amino acid residues at saturation equilibrium, which would prevent them from presenting as overlapped sites even though they are in fact saturated sites. This overlap ratio method is free of uncertain assumptions and hence more realistic than other methods of testing for saturation, such as comparing to inferred number of mutations based on uncertain phylogenetic trees derived from maximum parsimony or maximum likelihood methods or assuming saturation to mean fully random composition when in fact saturation (as a result of selection) could just as well mean a fraction of all possible choices at a site (Steel, Lockhart, Penny 1993; Philippe et al. 1994; Xia et al. 2003). Our findings here show that saturation is maintained by selection as fast evolving proteins have lower fraction of conservative changes, which corrects the presently popular view that variant sites at saturation are fully neutral.

By using the overlap ratio method, we have verified that the vast majority of proteins show maximum distances and only a small proportion, the slowest evolving, are still at the linear phase of changes (Huang 2010; Luo, Huang 2016; Yuan et al.2017). Variations at most genomic sites within human populations are also at optimum equilibrium as evidenced by the observation that a slight increase above the present genetic diversity level in normal subjects is associated with patient populations suffering from complex diseases (Yuan et al. 2012; Yuan et al. 2014; Zhu et al. 2015; Gui, Lei, Huang 2017; He et al. 2017; Lei, Huang 2017; Lei et al. 2018), as well as the observation that sharing of SNPs among different racial groups is an evolutionary rate dependent phenomenon with more sharing in fast evolving sequences (Yuan et al.2017).

It turns out that the original hemoglobin and cytochrome C alignment results of Zuckerkandl, Pauling, and Margoliash (Zuckerkandl, Pauling 1962; Margoliash 1963) also have high proportions of overlap sites and so represent in fact saturation distances (Huang 2010). Thus, the interpretation of those results as molecular clock or a linear distance phenomenon (mutations always increase distances and distances always correlate with time) was mistaken.

If fast evolving proteins are at saturation, they would not be expected to reveal authentic phylogenetic relationships. However, most past studies have used them and some meaningful results do emerge including the results of Zuckerkandl, Pauling, and Margoliash (Zuckerkandl, Pauling 1962; Margoliash 1963). So, how could saturation distances be correlated with time? It turns out that researchers have long overlooked the reality that similar sequences could also mean similar physiology. In a sequence alignment with humans, there is a hierarchy with humans less and less related to increasingly less complex species. As less complex species evolved earlier, the hierarchy of gene identities shows correlations with two different parameters, complexity and time. If one only focused on the time correlation as did Zuckerkandl, Pauling, and Margoliash, one would conclude that protein non-identity is only determined by time of separation as if the substitution rate is constant and the same among species (hence the molecular clock). Seemingly consistent with this, the same alignment data also showed that yeasts are equidistant to fishes, chickens, and humans, or that fishes are equidistant to chickens and humans (hence the genetic equidistance result). On the other hand, if one focused on the complexity parameter and ignored time, one would find a strong correlation of sequence identity with species complexity. One also finds that simple species is equidistant to all more complex species. So, the distance hierarchy with humans as measured by fast evolving proteins at saturation distance is a result of lower and lower complexity of species in more ancient times and hence higher and higher within species maximum genetic diversity (MGD). Lower complexity means wider tolerable error range in DNA building parts. The saturation distance to human for a lower complexity species is equal to the MGD of the lower species (Huang 2008a; Huang 2008b; Hu et al. 2013; Huang 2016).

The correlation of sequence identity with complexity makes sense as closer physiology should mean closer gene identity and less complex species should be able to tolerate more mutation variations (think HIV and bacteria). Genomes have two types of sequence mismatches, functional and neutral, both of which show correlation with time. The neutral variations are explained by the neutral theory. The functional variations are correlated with physiology, as explained by the maximum genetic diversity (MGD) theory, and indirectly with time as physiological complexity is correlated with time with simple physiology evolved earlier in time (Huang 2008a; Huang 2008b; Hu et al. 2013; Huang 2016). Functional variations are under physiological/natural selection and would quickly reach saturation maximum or optimum, because lower than optimum would mean less fitness such as poor immunity and so quick elimination. So, functional variations in functional DNAs are expected to be under positive selection to quickly reach optimum/maximum level, which explains why observed variants in fast evolving proteins are under more (positive) selection as found here.

Therefore, even though fast evolving proteins only represent maximum saturation distances, they may in some cases reveal proper phylogenetic topology due to the fact that complex species with lower MGD evolved more recently. Nonetheless, a proper method of phylogenetics must measure only ancestry-linked relationships and excludes sequence variations involved in physiology, which can only be accomplished by using slow evolving genes with fixed and standing neutral variations. The results here provided the key evidence for the neutrality of the fixed and standing variations in the slow evolving genes and should help with the popularization of using such genes in phylogenetic inferences.

Among the existing methods of phylogeny inferences, most, such as the maximum likelihood methods and Bayesian methods, require the assumption of certain evolutionary models of amino acid or nucleotide changes, which may be unrealistic (Felsenstein 1981; Rannala, Yang 1996). Distance matrix methods do not rely on such models but requires the molecular clock and hence the neutral variants. While maximum likelihood and related methods do not require the molecular clock, they do assume substitutions to be stochastic, probabilistic, and independent events (Felsenstein 1981), which only neutral variants can satisfy. As slow evolving proteins vary less dramatically in rates among species, they are inherently better suited for the distance matrix methods. We expect far more consistent future results of evolutionary trees to emerge once researchers have adopted a method that is largely free of uncertain assumptions.

## Acknowledgements

Supported by the National Natural Science Foundation of China grant 81171880 and the National Basic Research Program of China grant 2011CB51001 (S. H.).

## Declarations

### Ethics approval and consent to participate

Not applicable.

### Consent for publication

Not applicable.

### Availability of data and material

The datasets generated and analyzed for this study are available upon request.

### Author contributions

SH and MW designed the study. MW performed data analyses. DW and JY identified the slow evolving proteins. SH wrote the manuscript and all authors made comments and approved the final version.

## References

Auton, A, LD Brooks, RM Durbin, EP Garrison, HM Kang, JO Korbel, JL Marchini, S McCarthy, GA McVean, GR Abecasis. 2015. A global reference for human genetic variation. Nature 526:68–74.

Ayala, FJ. 1999. Molecular clock mirages. Bioessays 21:71–75.

Brandley, MC, Y Wang, X Guo, AN de Oca, M Feria-Ortiz, T Hikida, H Ota. 2011. Accommodating heterogenous rates of evolution in molecular divergence dating methods: an example using intercontinental dispersal of Plestiodon (Eumeces) lizards. Syst Biol 60:3–15.

Cai, JJ, DA Petrov. 2010. Relaxed purifying selection and possibly high rate of adaptation in primate lineage-specific genes. Genome Biol Evol 2:393–409.

Doolittle, RF, B Blombaeck. 1964. Amino-Acid Sequence Investigations Of Fibrinopeptides From Various Mammals: Evolutionary Implications. Nature 202:147–152.

Felsenstein, J. 1981. Evolutionary trees from DNA sequences: a maximum likelihood approach. J. Mol. Evol. 17:368–376.

Fitch, WM, E Margoliash. 1967. Construction of phylogenetic trees. Science 155:279–284.

Gui, Y, X Lei, S Huang. 2017. Collective effects of common SNPs and genetic risk prediction in type 1 diabetes. Clin Genet 93:1069–1074.

He, P, X Lei, D Yuan, Z Zhu, S Huang. 2017. Accumulation of minor alleles and risk prediction in schizophrenia. Sci Rep 7:11661.

Hu, T, M Long, D Yuan, Z Zhu, Y Huang, S Huang. 2013. The genetic equidistance result, misreading by the molecular clock and neutral theory and reinterpretation nearly half of a century later. Sci China Life Sci 56:254–261.

Huang, S. 2008a. Histone methylation and the initiation of cancer, Cancer Epigenetics. New York: CRC Press.

Huang, S. 2008b. Inverse relationship between genetic diversity and epigenetic complexity. Nature Precedings:doi.org/10.1038/npre.2009.1751.1032.

Huang, S. 2010. The overlap feature of the genetic equidistance result, a fundamental biological phenomenon overlooked for nearly half of a century. Biological Theory 5:40–52.

Huang, S. 2012. Primate phylogeny: molecular evidence for a pongid clade excluding humans and a prosimian clade containing tarsiers. Sci China Life Sci 55:709–725.

Huang, S. 2016. New thoughts on an old riddle: What determines genetic diversity within and between species? Genomics 108:3–10.

Kern, AD, MW Hahn. 2018. The Neutral Theory in Light of Natural Selection. Mol Biol Evol 35:1366–1371.

Kimura, M. 1983. The neutral theory of molecular evolution. Cambridge: Cambridge University Press.

Kimura, M, T Furukawa, T Abe, et al. 1998. Identification of two common regions of allelic loss in chromosome arm 12q in human pancreatic cancer. Cancer Research 58:2456–2460.

Kumar, S. 2005. Molecular clocks: four decades of evolution. Nat Rev Genet 6:654–662.

Lei, X, S Huang. 2017. Enrichment of minor allele of SNPs and genetic prediction of type 2 diabetes risk in British population. PLoS ONE 12:e0187644.

Lei, X, J Yuan, Z Zhu, S Huang. 2018. Collective effects of common SNPs and risk prediction in lung cancer. Heredity:doi:10.1038/s41437-41018-40063-41434.

Luo, D, S Huang. 2016. The genetic equidistance phenomenon at the proteomic level. Genomics 108:25–30.

Margoliash, E. 1963. Primary structure and evolution of cytochrome c. Proc. Natl. Acad. Sci. 50:672–679.

Pearson, WR. 2013. Selecting the Right Similarity-Scoring Matrix. Curr Protoc Bioinformatics 43:3 5 1–9.

Philippe, H, P Forterre. 1999. The rooting of the universal tree of life is not reliable. J Mol Evol 49:509–523.

Philippe, H, U Sorhannus, A Baroin, R Perass. 1994. Comparison of molecular and paleontological data in diatoms suggests a major gap in the fossil record. J. Evol. Biol. 7:247–265.

Pulquerio, MJ, RA Nichols. 2007. Dates from the molecular clock: how wrong can we be? Trends Ecol Evol 22:180–184.

Rannala, B, Z Yang. 1996. Probability distribution of molecular evolutionary trees: a new method of phylogenetic inference. J Mol Evol 43:304–311.

Soubrier, J, M Steel, MS Lee, C Der Sarkissian, S Guindon, SY Ho, A Cooper. 2012. The influence of rate heterogeneity among sites on the time dependence of molecular rates. Mol Biol Evol 29:3345–3358.

Steel, MA, PJ Lockhart, D Penny. 1993. Confidence in evolutionary trees from biological sequence data. Nature 364:440–442.

Xia, X, Z Xie, M Salemi, L Chen, Y Wang. 2003. An index of substitution saturation and its application. Mol Phylogenet Evol 26:1–7.

Yuan, D, X Lei, Y Gui, Z Zhu, D Wang, J Yu, S Huang. 2017. Modern human origins: multiregional evolution of autosomes and East Asia origin of Y and mtDNA. bioRxiv:doi: >https://doi.org/10.1101/106864.

Yuan, D, Z Zhu, X Tan, et al. 2012. Minor alleles of common SNPs quantitatively affect traits/diseases and are under both positive and negative selection. arXiv:1209.2911.

Yuan, D, Z Zhu, X Tan, et al. 2014. Scoring the collective effects of SNPs: association of minor alleles with complex traits in model organisms. Sci China Life Sci 57:876–888.

Zhu, Z, X Man, M Xia, Y Huang, D Yuan, S Huang. 2015. Collective effects of SNPs on transgenerational inheritance in Caenorhabditis elegans and budding yeast. Genomics 106:23–29.

Zuckerkandl, E, L Pauling. 1962. Molecular disease, evolution, and genetic heterogeneity, Horizons in Biochemistry. New York: Academic Press.

